# Electrical information flows across the sporocarps of two ectomycorrhizal fungi in the field

**DOI:** 10.1101/2025.09.01.673447

**Authors:** Yu Fukasawa, Daisuke Akai, Takayuki Takehi, Daiki Takahashi, Yutaka Osada

## Abstract

We measured the extracellular bioelectrical activities of two ectomycorrhizal basidiomycete species belonging to the genus *Hebeloma* in a field to examine the potential information flow across the sporocarps and its reponse to artificial stimulation. The 37 sporocarps (29 *H. danicum* and 8 *H. cylindrosporum*) occurring within a 5 m × 5 m quadrat on the forest floor of an oak-dominated stand were measured for their electrical potential every second for 3.5 days using subdermal stainless electrodes. Causality analysis of electrical potential during a period without artificial stimulation indicated that the magnitude of potential information flow across the sporocarps was not restricted to the sporocarps belonging to the same genet (clone) nor within a species. Howeverm this was still negatively associated with genetic as well as spatial distances. The effects of artificial stimulation (water and urine addition) on the average magnitude of information flow across the sporocarps were positive when tap water was added locally to a certain sporocarp but were negative when tap water was added to all the sporocarps. In contrast, the addition of urine had minimal effect on the magnitude of information flow. These results indicate the presence of underground electrical information flows across fungal sporocarps which respond vigorously to the environmental conditions.

## Introduction

Fungi develop vast mycelial networks in the litter-soil interface of the forest floor and have an important role in nutrient and carbon dynamics by decomposing dead plant materials [1] and transporting water, nutrients, and carbon across their network [2]. Recent studies suggest that mycorrhizal fungi networks connect multiple plants and transport materials as well as signals between mycelia and plants under laboratory conditions [3]. However, such signal transfer through fungal mycelia has rarely been recorded in the field [4].

Bioelectricity is used for signal transfer in various organisms, from brained animals to single-celled bacteria [5]. In fungi, electricity has been measured in single hyphae [6], mycelium [7], and sporocarps [8]. Adamatzky [8] reported that electricity patterns caused by artificial stimulation, such as flame and salt, could be transported across sporocarps of cultivated oyster mushrooms. Under field conditions, Fukasawa et al. [9] recorded patterns of electrical potentials of six sporocarps of the ectomycorrhizal fungus *Laccaria bicolor* on the forest floor using a subdermal electrode and indicated that electricity patterns might be transported across sporocarps with directionality. However, this preliminary data was only measured on a limited number of sporocarps within a short distance. Further research is required on a larger spatial scale to investigate the general features of electrical signal transfer across the fungal mycelial network in the field.

Fungi are microorganisms, and their individual hyphae are approximately 10 µm thick and invisible to the naked eye, and thus it is challenging to record their electrical potential in the field. In this regard, sporocarps emerging on the ground surface are large and strong enough to be settled with electrodes for a while and are thus a good option to measure the electrical potential [8]. However, the occurrence of sporocarps is highly dependent on environmental conditions, such as temperature and precipitation, and therefore, the opportunities to measure the electrical potential of multiple sporocarps are limited by chance. To overcome this limitation, we focused on ammonia fungi. Ammonia fungi are an ecological group of fungi that prefer and fruit in the environmental conditions created after the addition of high concentrations of ammonia [10]. By adding urea to the forest floor in spring as an ammonia precursor, we can expect mass fruiting of ectomycorrhizal ammonia fungi in the urea-fertilized area in the fall.

In the present study, we conducted field experiments to stimulate genetically identified ammonia fungi (genetically identical clones, possibly connected by hyphae underground) and investigate the effects of genetic and spatial distance on electrical activities and information flow between the sporocarps. This experiment became possible for the first time with the introduction of the urea-induced mass fruiting technique. We hypothesized that the spatial distance between sporocarps has a significant effect on the magnitude of electrical information flow between the sporocarps because a previous study showed that spatially closer sporocarps exhibited greater electrical information flow [9]. Moreover, we hypothesized that the genetic distance between the sporocarps has a significant effect on the magnitude of electrical information flow between the sporocarps, because mycelial connectivity might be important for the transduction of electrical activity. We predicted stronger information flow within the identical genet compared to that of genetically distant sporocarps of the same species or different species that may not be connected by the mycelial network. Our third hypothesis is that different stimulations induce different electrical information flow of the sporocarps because different stimulants induce different electrical responses in fungi [8]. We investigated the addition of tap water and urine as pulse stimulants. We predict that the electrical response of the sporocarps is greater when urine is added compared to that of tap water, because the focal sporocarps are ammonia fungi, and urine is an ammonia precursor in soil.

## Materials and methods

### Study site

This study was performed in a secondary mixed forest dominated by *Quercus serrata* in Kami town, Miyagi, Japan (38°37′23.29″ N, 140°48′57.53″ E, 130 m a.s.l.). In 2022, the mean annual temperature at the nearest meteorological station at Kawatabi (38°45′19.28″ N, 140°45′14.38″ E, 321 m a.s.l.) was 11.1°C (range: –1.2°C in January to 23.6°C in July), and the annual precipitation was 1675.5 mm. Snow covers the ground from December to March (Japan Meteorological Agency, https://www.jma.go.jp/jma/indexe.html). On April 18, 2022, a 5 m × 5 m study plot was established in the study site, and urea (640 g m^−2^) was added to three spatially separated areas in the plot to stimulate ammonia fungi fruiting (Fig. 1A).

**Fig. 1.**
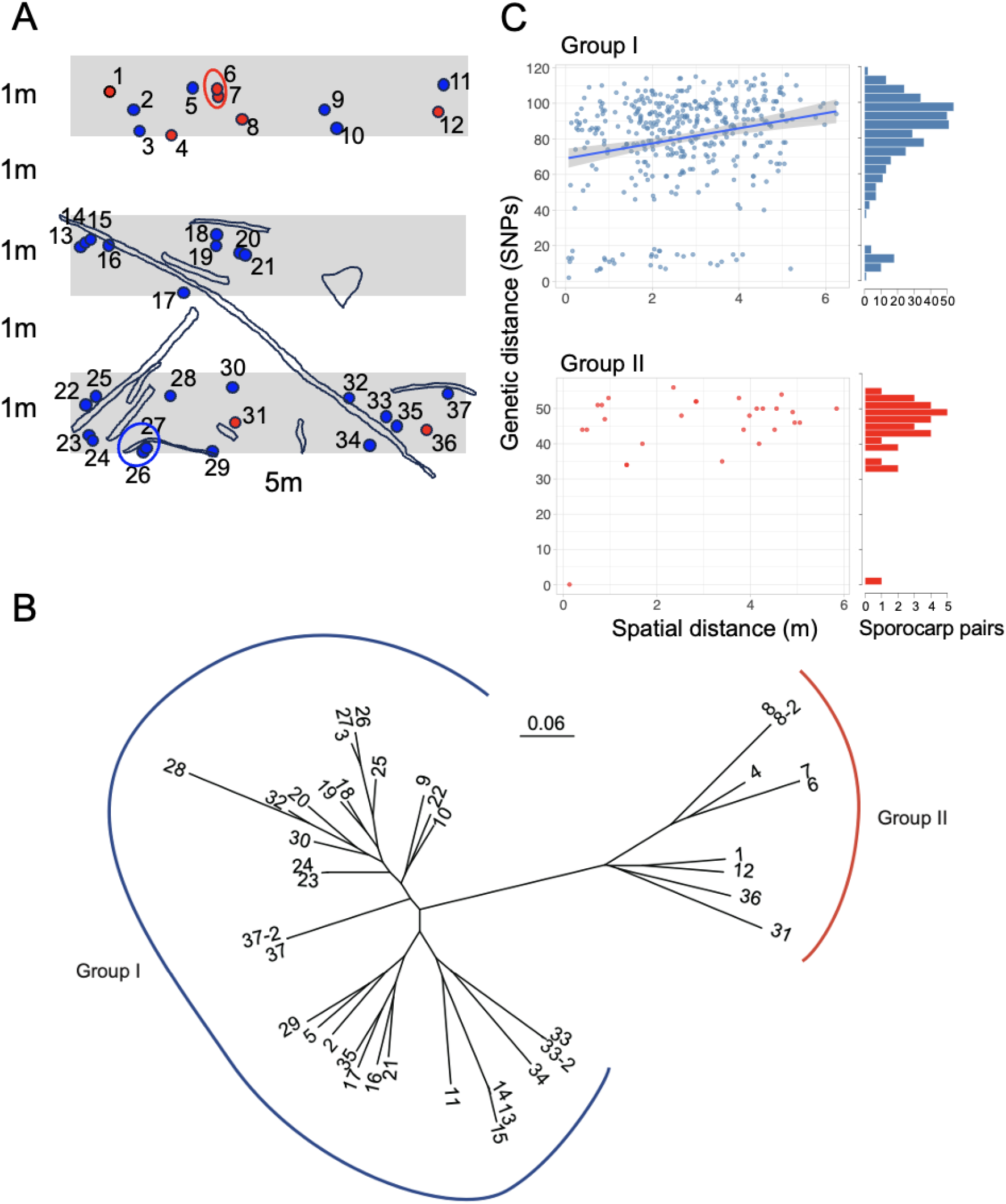
(A) Map of the study plot on October 23, 2022, showing locations of sporocarps#1– #37. Urea was added to shaded areas on April 18 to stimulate sporocarp fruiting of ammonia fungi. Rectangular shapes are fallen branches of *Quercus serrata*. A triangle-like shape at the middle is the base of a *Q. serrata* snag. Blue and red dots represent sporocarps of Groups I and II, respectively. Dots surrounded by blue and red circles indicate clusters of genetically identical sporocarps (genets) of the Groups I and II, respectively. (B) Phylogenetic relationships among sporocarps#1–#37. The maximum likelihood tree was constructed from 12,073 SNPs obtained via MIG-seq analysis. Sporocarp#8 and #8-2, #33 and #33-2, and #37 and #37-2 represent samples and their duplicates, respectively. (C) Scatter plot of the sporocarp’s pairwise spatial and genetic distances, with histograms showing the distribution of genetic distance in Groups I and II. A regression line with a 95% confidence interval was shown in Group I because the correlation was statistically significant (Pearson’s *r* = 0.226, *p* < 0.001) but was not significant in Group II.

### Recording the electrical activities of fruit bodies

On October 23, 2022, we found multiple fruitings of sporocarps in the plot (Fig. 1A, Fig. 2). Some occurred within a cluster, and others formed discrete groups. We inserted electrodes into the cap and stipe for 37 sporocarps (sporocarp#1–#37), with the recording electrode on the cap and the reference electrode on the stipe, to record the electrical potential difference between the cap and stipe (Fig. 2). We used subdermal needle electrodes (SPES MEDICAL SRL Via Buccari 21 16153 Genova, Italy). The distance between the recording and reference electrodes in a fruit body was approximately 4 cm. Recording started at 11:45 on October 23 (Day 1) and stopped at 8:00 on October 27 (Day 5). Electrical activity of the sporocarp was recorded using a GRAPHTEC GL840-M midi logger (GRAPHTEC, Kanagawa, Japan) with a voltage range of –100 to 100 mV. Electrical activity was recorded every second.

**Fig. 2.**
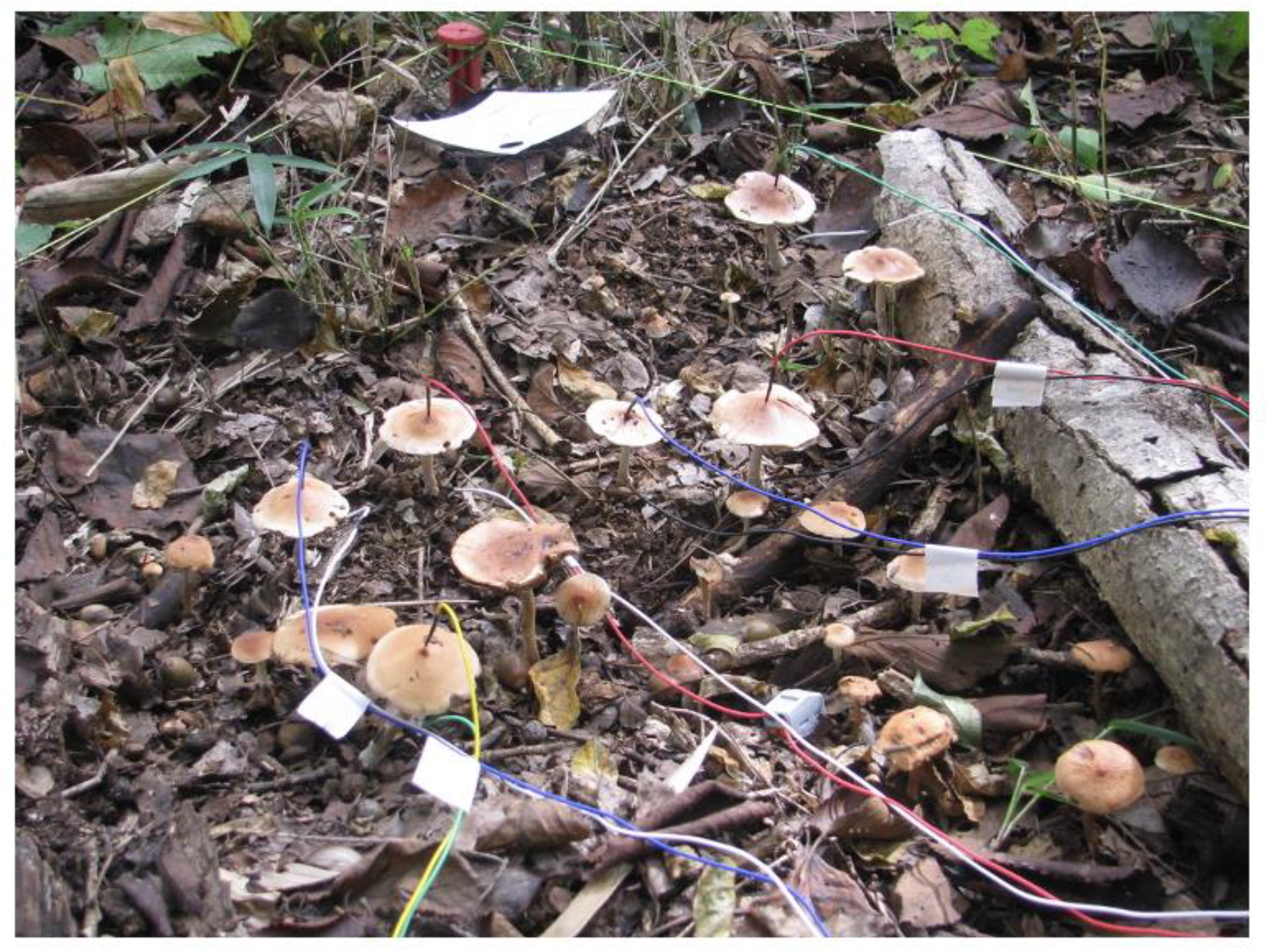
Sporocarps with electrodes. The recording electrode was inserted on the top of the cap, and the reference electrode at the bottom of the stipe.

During recording, we performed stimulation experiments by adding 200 mL of tap water or urine to the bottom of sporocarp#1 at 20:31 on Day 1 (Stim0, water), 5:52 on Day 2 (Stim1, urine), 5:55 on Day3 (Stim2, water), and 6:47 (Stim4, urine) and 17:13 on Day 4 (Stim5, urine). The urine was collected from one of the authors (YF, 44-year-old male without any chronic disease). In addition, tap water (10 L m^−2^) was added to all the area treated with urea (shaded area in Fig. 1A) to maintain the sporocarp’s electrical activities at 5:55 on October 25 (Stim3). The water and urine were gently added to the ground without spraying directly onto the sporocarps and electrodes.

The daily mean temperature at Kawatabi, the nearest meteorological station, was 12.3°C on Day 1, while it decreased to 7.6°C on Day 2 and continued decreasing to 6.6°C on Day 5 (Japan Meteorological Agency, https://www.jma.go.jp/jma/indexe.html). The study site experienced little rain during recording except for a short shower on Day 1, which was not sufficient to be recorded by the meteorological system.

### Clonal and species identification of sporocarps

At the end of recording on October 27, the 37 sporocarps were collected, brought to the laboratory, and stored at –30°C until DNA extraction using the CTAB method [11]. Clonal analysis was performed by using genome-wide SNPs obtained from multiplexed inter-simple sequence repeats genotyping by sequencing (MIG-seq) analysis [12,13]. To distinguish true genetic differences among different genets from potential PCR and sequence errors, we included sample duplicates (sporocarps#8, #31, and #37). Library preparation for the MIG-seq analysis was performed as described by Suyama et al. [13]. Sequencing was performed on the Illumina MiSeq platform (Illumina, San Diego, CA, USA) by using the MiSeq Reagent Kit v3 (150-cycle) (Illumina). The data were deposited in the National Center for Biotechnology Information (NCBI; BioProject ID: PRJNA1165506) database. Forward and reverse data of each sample were merged and analyzed as single-end data following Suyama and Matsuki [12]. Raw reads were filtered and trimmed using Trimmomatic v.0.32 software [14] with the following options: HEADCRAP:6, CROP:77, SLIDINGWINDOW:10:30, and MINLEN:51. We assembled datasets using the Stacks v.2.41 pipeline [15], and default values were employed for other parameters.

We performed phylogenetic analysis using all samples and divided the datasets based on their genetic relationships to include as many SNPs as possible in clonal analysis. Phylogenetic analysis was performed using a maximum likelihood search with the GTR+G model using raxMLGUI v.2.0 [16]. We extracted SNPs with a minimum genotyping rate of 10% (-R 0.1) using populations software of Stacks pipeline, and 12,073 SNPs were used for phylogenetic analysis. According to the result (Fig. 1B), the samples were divided into two groups (Group I: 31 samples including two duplicate samples, and Group II: 9 samples, including one duplicated sample). We extracted SNPs with a minimum genotyping rate of 90% (-R 0.9), and clonal analysis was performed by using 360 and 87 SNPs for Groups I and II, respectively. For each dataset, we calculated pairwise genetic distance (number of SNPs) between samples, and samples with a distance below the threshold value were assigned to the same genet. The threshold value for each dataset was set by the number of SNPs between the duplicated samples (four in Group I, and two in Group II). Clonal analysis was performed using GenoDive version 3.05 [17].

DNA samples of the three sporocarps were randomly selected from each of the two groups (sporocarp#9, #21, and #34 for Group I; #6, #8, and #12 for Group II) and identified by clonal analysis, and their internal transcribed spacer (ITS) region was analyzed further for species identification. Fungal rDNA was PCR-amplified using the ITS1F and ITS4 primer pair [18]. The PCR products were purified using Multiplex PCR Assay Kit Ver. 2 (Takara Bio, Kusatsu, Japan). Sequencing was performed by FASMAC (Atugi, Japan) using the same primer pair. An ITS region with a 97% shared identity was used as the criterion for the molecular identification of species [19]. BLAST searches were performed against publicly available sequences in the NCBI database (accession#: LC844115 for Group I; LC844116 for Group II) to identify the sporocarp species.

### Causality analysis of electrical potentials

Using the time-series data of electrical activity from the 37 sporocarps, we estimated transfer entropy (*TE*, [20]) to identify potential causal relationships among the sporocarps. Transfer entropy, a generalization of Granger causality, quantifies the directional flow of information between two entities in a nonparametric manner [21]. More precisely, it defines the flow of information from a causal variable (*y*_*t*−*p*_) to an effect variable (*x*_*t*_) with a causal time delay *p*, as follows:

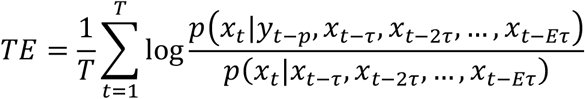

where *E* and *τ* represent the embedding dimension of the effect variable and time interval required for the embedding time delay, respectively, and *T* is the number of time points used to calculate *TE*. The unified information-theoretic causality (UIC) method was used to estimate *TE* and assess its significance [22]. To reduce bias effects because of the small sample size, we applied effective transfer entropy (*ETE*), which adjusts for *TE* bias using surrogate data [23].

Before performing UIC analysis, the time-series data of the electrical activity were differenced to create a stationary time-series. Moreover, we separated the data into several time windows for analysis (Fig. 3). The first and second windows were each 30 min of data without artificial stimulation (resting periods 1 and 2). A 45-min break was inserted between resting periods 1 and 2, accounting for the brief rain shower. The successive windows are each 9 h for the six stimulation events described above (Stim0–5, addition of water or urine). For each window, the stimulation event occurred in the middle of the 9 h data window, consisting of 4.5 h before and after each event. Each 9-h data set was divided into 18 segments (30 min each). The embedding dimension (*E*) was selected to enhance the predictive accuracy of the simplex projection method [24]. The time interval (*τ*) was set to the recording interval (i.e., 1 s) because the electrical activity of the sporocarps is expected to change within a shorter time scale than the recorded interval. In the UIC analysis, we calculated the *ETE* of aggregated data across the entire record of each time segment, which enabled us to examine the average trends of the information flow in each segment. The analysis was conducted using a causal time delay (*tp*) of –30 to 0 s with a 1 s interval to assess the duration of causal influence. We calculated the maximum *ETE* and associated *tp* (–30 to 0 s) in each time segment for subsequent analyses.

**Fig. 3.**
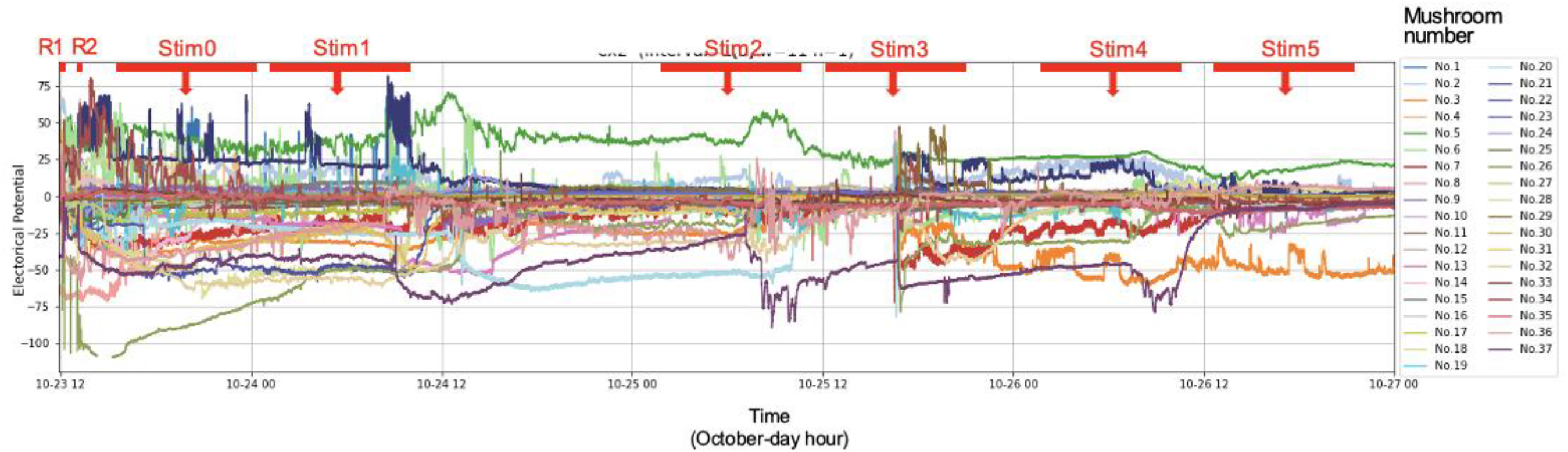
Full data of the sporocarp’s electrical potentials and position of time windows used for causality analysis (red bars). A 30-min window for resting period 1 before rain (R1), a 30-min window for resting period 2 after rain (R2), and a 9 h window for each of Stim#0–5, with 4.5 h for before and after the stimulation timing, which are shown as red arrows in the figure.

## Statistical analysis

All analyses were performed using R version 4.4.2 [25]. Pearson’s correlation coefficient was calculated between spatial and genetic distances of the sporocarps for each of the Group I and II datasets. The *ETE* matrix of the sporocarps during resting period 1 was used to visualize the network of information flow across all 37 sporocarps. The *ETE* and *tp* matrices during resting period 1 were used to analyze the relationships between effect *ETE* and causal *ETE*, as well as between effect *tp* and causal *tp* of the 37 sporocarps, which were visualized in a two-dimensional space and as a cluster dendrogram using the Ward method. The shift of the relationship between effect *ETE* and causal *ETE*, and between effect *tp* and causal *tp* of the 37 sporocarps during resting periods 1 and 2, was also shown in two-dimensional space.

The Mantel test was used to analyze relationships between causality indices and the sporocarps’ spatial and genetic distances using the *mantel* command in the *vegan* package [26]. Mantel tests were performed for each Group I and Group II dataset and the combined dataset. For the combined dataset, species identity was used as the genetic distance, whereas the number of SNPs was used for each group dataset. The *ETE* and *tp* asymmetric matrices were converted to be symmetric by taking the averages of the opposite matrices, i.e., (*ETE*_ij_ + *ETE*_ji_) /2, where i and j are sporocarp individuals. Furthermore, to make the correlation between causality indices and spatial and genetic distances positive, the reciprocal of causality indices was used for Mantel correlation analysis.

The effects of artificial stimulation events on information flow across the sporocarps were evaluated using the Regression Discontinuity Design (*rdrobust* command in the *rdrobust* package). The *ETE* across the 37 sporocarps at each of the 18 time segments was compared before and after the stimulation event by setting the threshold between time segments 8 and 9 (the timing of the stimulation event).

## Results

### Genetic relationship of the sporocarps

Amplicon sequencing of the rDNA ITS region indicated that three sporocarps from Group I (#9, #21, and #34) and three from Group II (#6, #8, and #12) exhibited 100% similarity within each group, respectively. The BLAST result of the ITS sequence of Group I showed 99.85% similarity with *Hebeloma danicum* (accession# MT117079.1), and that of Group II revealed 98.38% similarity with *H. cylindrosporum* (accession# FJ769363.1). These ITS sequences are provided in supplementary FASTA file.

MIG-seq showed that sequences of the sporocarp#33 and #37 in Group I had four SNPs (genetic distance) between their duplicate samples (Supplementary Table S1) and two SNPs in the duplicate samples of sporocarp#8 in Group II (Supplementary Table S2). There were <4 SNPs between sporocarp#26 and #27 in Group I and thus regarded as an identical genet (Table S1; Fig. 1A). Similarly, there were <2 SNPs between sporocarp#6 and #7 in Group II and were also regarded as another identical genet (Table S2; Fig. 1A). In Fig. 1C, a histogram of full-factorial pairwise SNPs across the sporocarps revealed bimodal distributions in both groups, with 20–30 SNPs as the gap between the peaks. In Group I, sporocarp#28 showed <20 SNPs with all other sporocarps (Supplementary Table S1). Similarly, sporocarp#13, #14, and #15, and sporocarp#23 and #24 showed <20 SNPs with each other, respectively. There was a significant correlation between spatial distance and SNPs among the sporocarps in Group I (*r* = 0.226, *p* < 0.001), whereas no correlation was observed in Group II (Fig. 1C).

### Electrical potential

The electrical potential of the sporocarps fluctuated from the start of recording, ranging from –110 to 80 (Fig. 3, Supplementary Fig. S1 for data of each sporocarp). Most of the sporocarps responded to the stimulation event Stim3, which was water added to the areas that received urea. Other stimulation events, except for Stim2, also altered some electrical potential of sporocarp#1, but their effects on other sporocarps were not clear.

*ETE* of the electricity data during resting period 1 before rain revealed the network of information flow across 37 sporocarps, where sporocarp#1 exhibit the strongest information flow to the distant sporocarp#23 of a different species (Fig. 4). Similarly, sporocarp#1 tended to have strong information flow toward sporocarps #2, #3, #5, and #15. Relationships between causal and effect *ETE* for the 37 sporocarps are shown in Fig. 5A, which were categorized into two clusters (Fig. 5C). One consists of the sporocarps showing larger effect *ETE* than causal *ETE* (dots on the left above the y = x line), and the other showing larger causal *ETE* than effect *ETE* (dots on the right below the y = x line). The species identity of sporocarps was not associated with this clustering. Moreover, the same relationship was shown between effect and causal *tp* value (Fig. 5B, D). Sporocarp#1 showed relatively large causal *ETE* but relatively small effect *ETE*, as well as relatively large causal *tp* and small effect *tp*. However, the clustering of *ETE* and *tp* was not always the same. For instance, sporocarp#1 and #31 were in the same *ETE* category (Fig. 5C) but in different *tp* categories (Fig. 5D). A weak but significant correlation was detected between *ETE* and *tp* (Supplementary Fig. S2) (*r* = 0.0538, *p* = 0.04963). After the rain between resting periods 1 and 2, the relationships between causal and effect *ETE* and between causal and effect *tp* changed substantially for most of the sporocarps (Fig. 6).

**Fig. 4.**
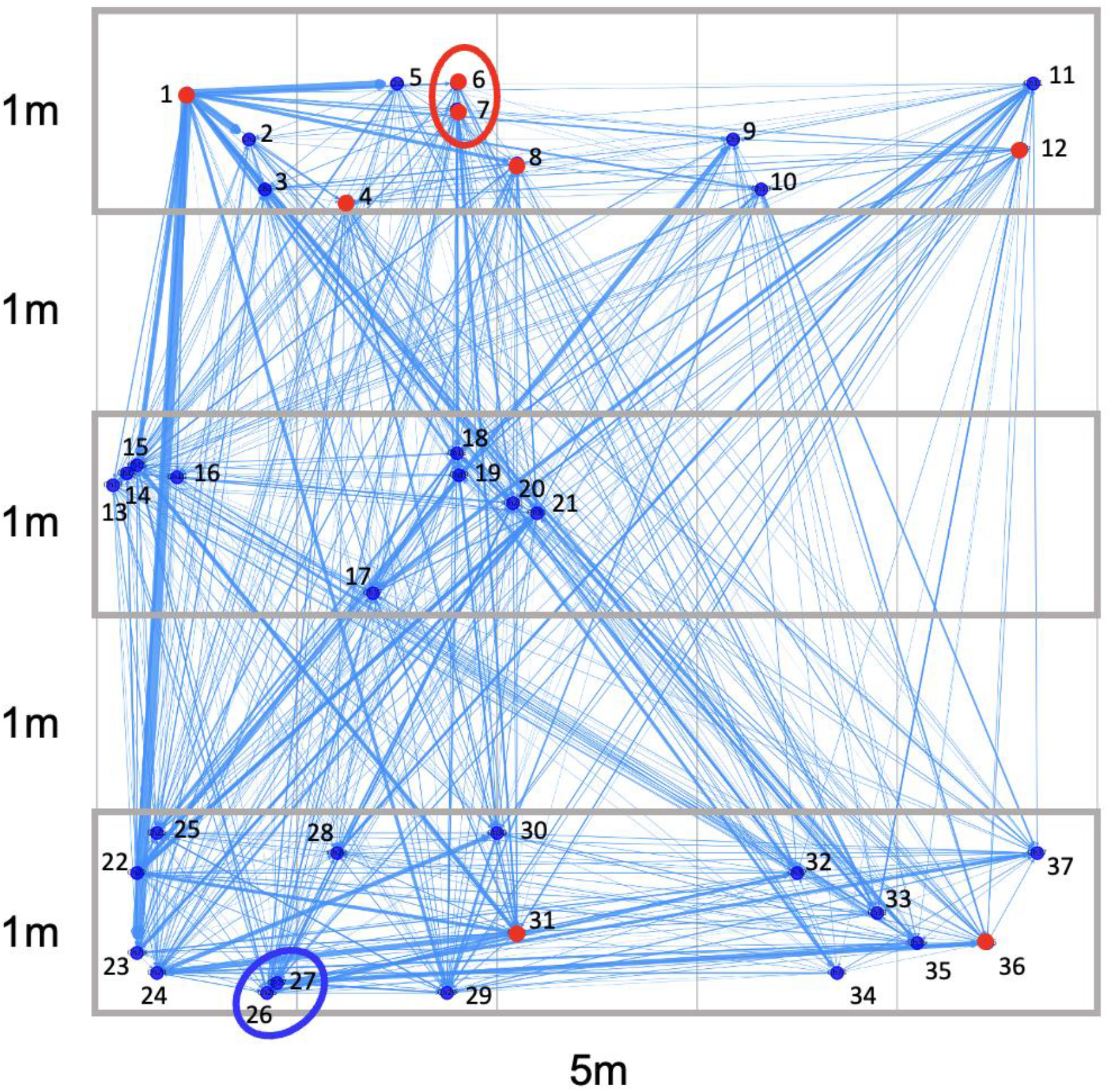
Network of information flow across sporocarps in the study plot during resting period 1. The thickness of blue arrows represents the magnitude of information flow (*ETE*). Only significant (*p* < 0.05) flows were shown. Dots show the location of investigated sporocarps: blue, Group I; red, Group II. Blue and red circles enclose genetically identical sporocarp pairs in Groups I and II, respectively. The area enclosed by gray squares are the area where urea was added in the spring.

**Fig. 5.**
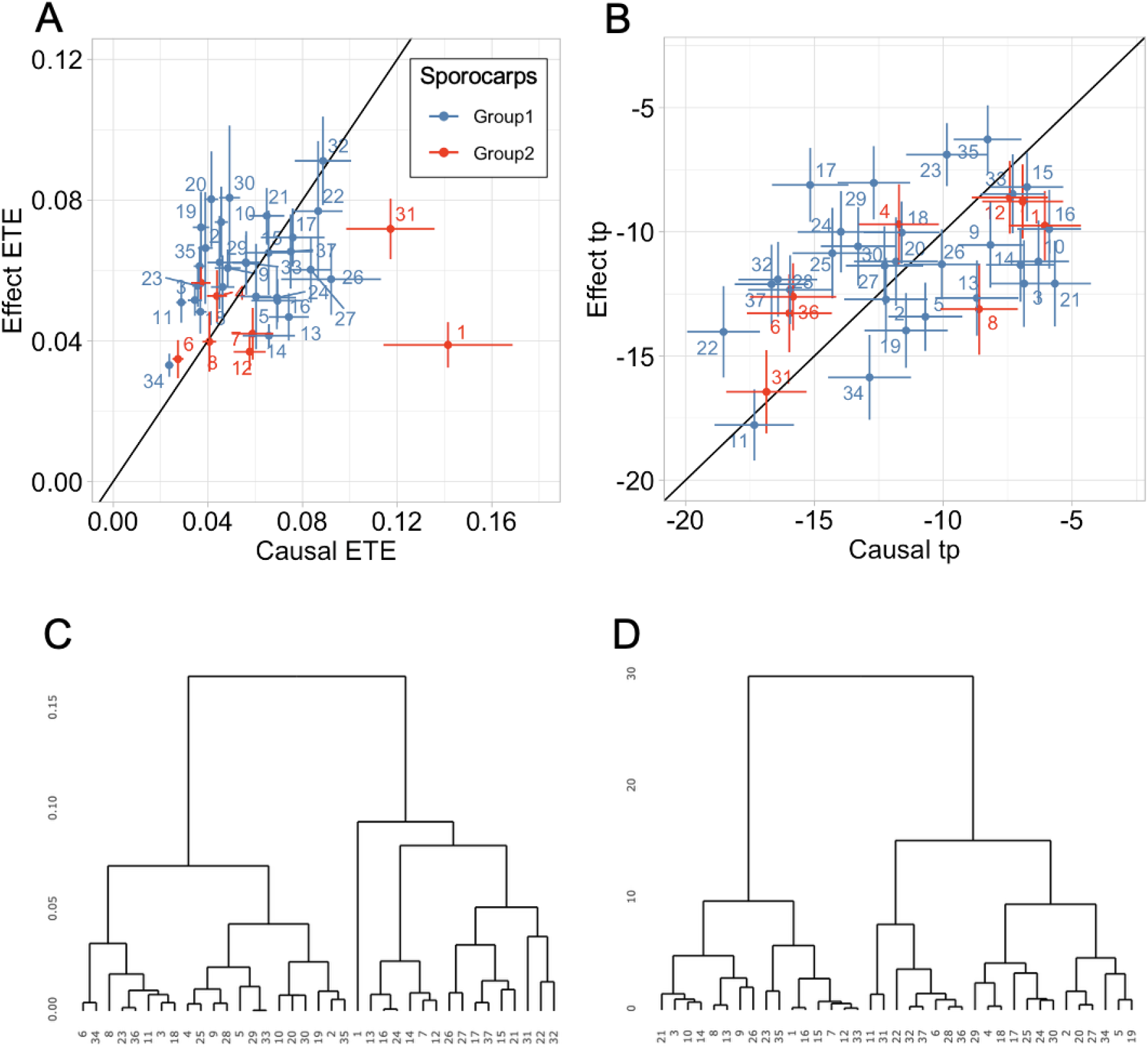
Relationship between causal and effect *ETE* (A) and *tp* (B) in the 37 sporocarps during resting period 1. Dots and error bars represent the mean and standard errors of 36 causal relationships with other sporocarps. Slopes in the figures indicate the y = x lines. Cluster dendrogram of the 37 sporocarps drawn using the Ward method by the distance calculated from causal and effect *ETE* (C) and *tp* (D).

**Fig. 6.**
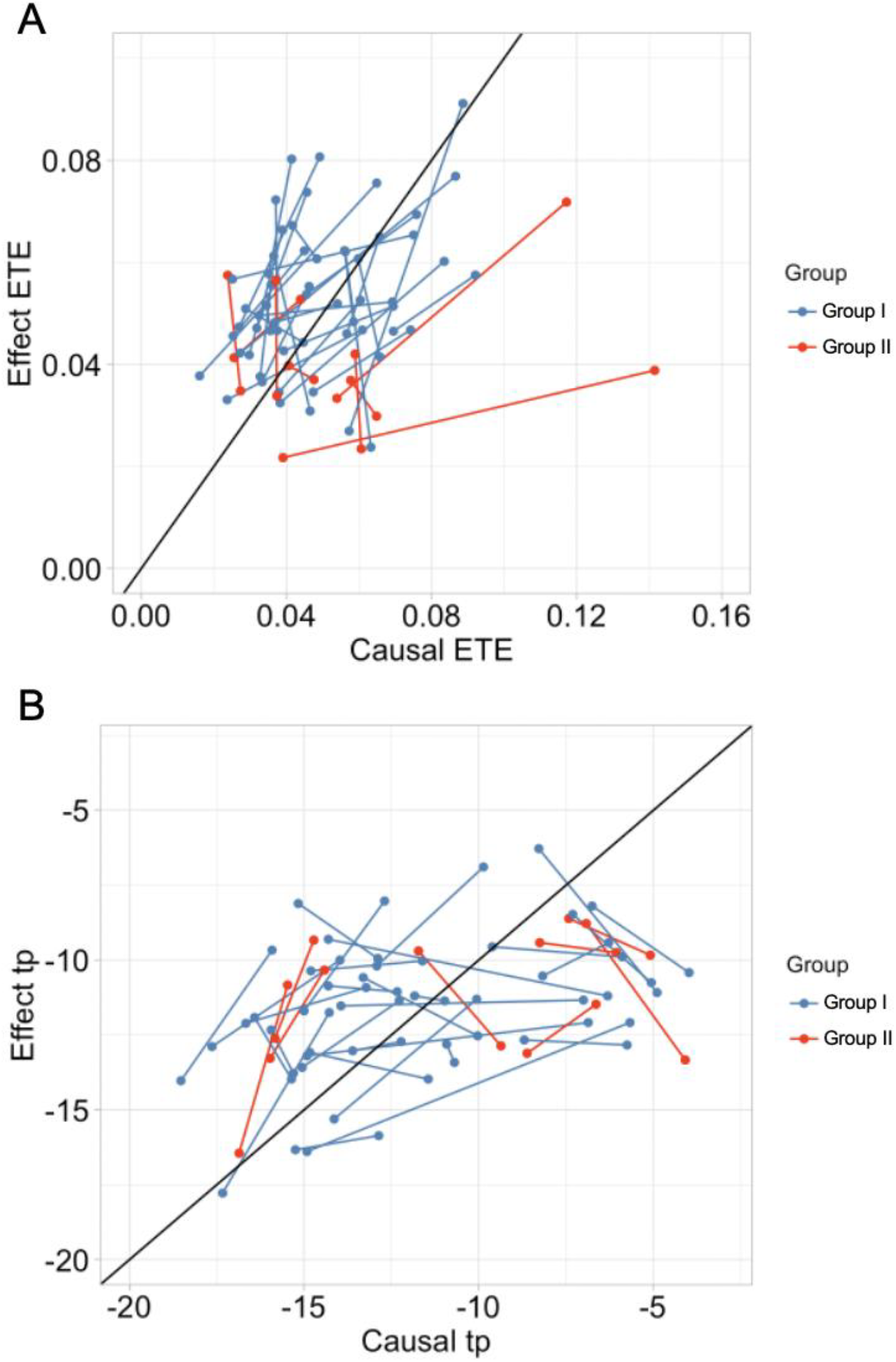
Relationship between causal and effect *ETE* (A) and *tp* (B) in the 37 sporocarps during resting period 1 (before rain) and resting period 2 (after rain), connected by a line. The identity of the resting period (1 or 2) was not identified in the figure. The black slopes in the figures indicate the y = x lines. The data for resting period 1 are the same as Fig. 5, and error bars are not shown for the sake of simplicity.

The Mantel test for the combined dataset showed that the spatial distance (Mantel’s *r* = 0.1141, *p* = 0.014) and species identity (Mantel’s *r* = 0.1386, *p* = 0.028) were positively associated with the *ETE* reciprocal. For the group datasets, the reciprocal of Group I *ETE* was significantly (Mantel’s *r* = 0.1097, *p* = 0.033) associated with the sporocarp’s spatial distance but not with their genetic distance. Neither the spatial nor the genetic distances of Group II sporocarps were significantly associated with *ETE*. The reciprocals of *tp* had no associations with spatial distance and species difference (and genetic distance).

Fig. 7 shows the changes in the mean *ETE* of the information flow across the 37 sporocarps during time segments, including the six artificial stimulations. The changes were dependent on the type of disturbance. Stim1, Stim4, and Stim5, where 200 mL of urine was added to the ground near the sporocarp#1, did not change the mean *ETE* of the 37 sporocarps, except for a small but significant change after Stim1. However, addition of 200 ml tap water to the ground near the sporocarp#1 (Stim0 and Stim2) significantly increased the mean *ETE*, and the addition of tap water onto the whole area treated with urea (Stim3) greatly reduced the mean *ETE* of the 37 sporocarps.

**Fig. 7.**
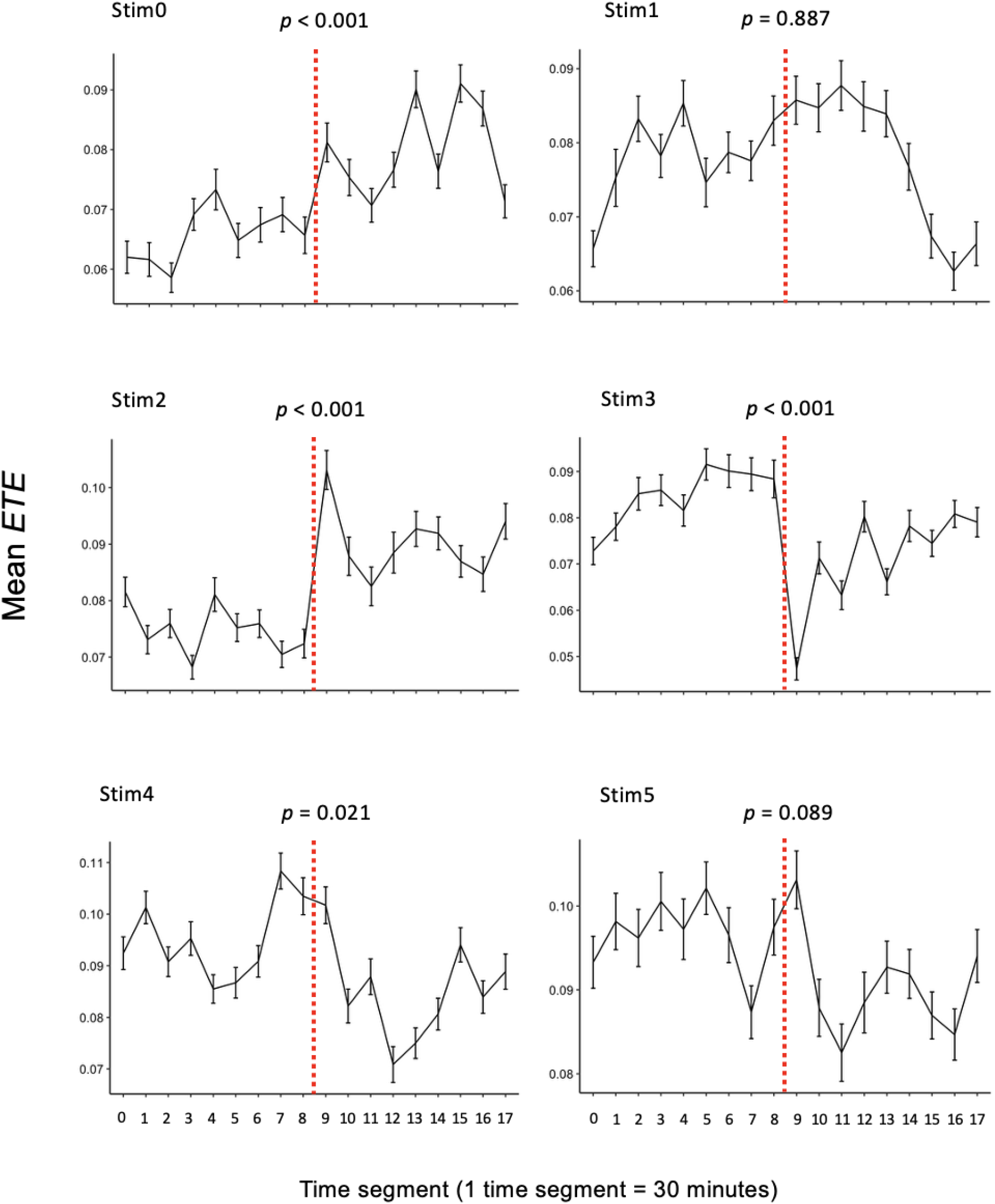
Mean *ETE* at each time segment for 4.5 h before and after the five stimulation events (totally 9 h for each window). Stim0 and Stim2: 200 mL of water added to sporocarp#1; Stim1, Stim4, and Stim5: 200 mL of urine to sporocarp#1; Stim3: tap water (10 L m^−2^) added to the whole area treated with urea. Error bars show the standard error. The effect of stimulation events between time segments 8 and 9 (shown as a dotted line) was evaluated using the Regression Discontinuity Design (*p*-values were shown in the figure).

## Discussion

In contrast to a previous report [9], where little electrical potential was detected at the beginning, the sporocarps’ electrical potential was detected from the start of the recording in the present study. In addition to the potential effect of different fungal species (*L. bicolor* in the previous paper), precipitation timing may be a reason for this difference. In the previous study, the sporocarps experienced little precipitation for two weeks before recording. Whereas, in this study, the sporocarps experienced rain (4 mm) a day before the start of recording (Japan Meteorological Agency), which could activate the sporocarps’ electrical potential [9]. After the start of recording, little rain was experienced except for a small rain shower between resting periods 1 and 2. Therefore, the fluctuations in electrical potentials recorded during resting period 1 might not be due to the direct effects of raindrops (inherent electrical charges, nutrient ions, and physical impacts [27–29]). Several abiotic and biotic factors could affect the electrical potential of the sporocarps. For example, flickering sunlight filtered through trees may affect the electrical potential of fungal sporocarps, as light can alter the electricity of fungi [30,31]. Various organisms, such as slugs and insects, eat fungal sporocarps by grazing [32,33], and such physical disturbance could also alter the electricity of fungi by increasing Ca^2+^ concentration in hyphal cells and generating membrane potential [34].

*ETE* was calculated using time-series data of sporocarps’ electrical potentials and indicates the potential information flow across the sporocarps [22]. In the present study, information flow across the sporocarps was analyzed on the largest scale to date (in terms of spatial and sporocarp number) using causality analysis of time-series data of electrical potentials. The *ETE* of the 37 sporocarps during resting period 1 without artificial stimulation before rain indicated that substantial information flows was detected between sporocarps of the same and different species. The mechanism of this interspecific information flow was not clear. A potential mechanism is chemical signaling across species. Adjacent fungal colonies interact with each other by secreting various chemicals [35–37], which may affect the physiology of neighboring colonies of different fungal species [38] and alter their electrical potential. Given that the sporocarps are made of interwoven hyphae, the electrode inserted into the caps and stipes measured the extracellular electrical potential difference between the cap and stipe. Such potentials could be influenced by not only membrane potential of fungal cells but also pH conditions altered by the secretion of organic acids, which can change the extracellular ionic balance [39–41]. This is expected to affect the electrophysiology of neighboring fungal colonies regardless of species.

The Mantel test revealed that spatial distance has a significant effect on information flow across sporocarps. The stronger information flow in the closer location is consistent with a previous study [9]. In addition, the present study showed that genetic distance within a species, as well as species difference, has a significant effect on the magnitude of information flow across sporocarps, which is a novel finding. However, genetically identical sporocarps did not show particularly stronger information flow with each other compared to that of sporocarps of the same species but genetically distant and of different species. Two explanations are possible for this phenomenon. First, the information flow across the sporocarps might not be through hyphae, and thus, the direct hyphal connection between clones is not necessary. In contrast, the second explanation is based on the speculation that most of the sporocarps (particularly in Group I) were connected by underground hyphae instead of their genetic distances. Previous studies reported that the heterothallic nature of fungi induces genetic variation across cells even within a single mycelium [42–44]. The histogram of SNPs in sporocarps of both groups was bimodal (Fig. 1C), suggesting that sporocarps included in the fewer SNPs peak were genetically closer than those in the other peak. In the case of sporocarps in Group I (identified as *H. danicum*), the largest number of SNPs in the fewer SNPs peak was 18, which was larger than 4, defined as the threshold of clones based on the duplicate samples in the present study. However, despite the relatively large number of SNPs, these genetically distant sporocarps could be included in the same mycelial network as discussed above. If so, sporocarp#28 might be connected to the other 28 sporocarps in Group I, and thus all 29 sporocarps in Group I were connected by a mycelial network. The Mantel test using data of Group I sporocarps indicated that only spatial distance was significantly associated with *ETE*, and genetic distance had no effect. This result supports the speculation that all sporocarps in Group I were connected to the same mycelial network. Although few studies have shown the size of underground mycelial network of *Hebeloma* spp., those of ectomycorrhizal *Suillus bovinus, S. pictus*, and *Rhizopogon vesiculosus* were ≥20 m, and that of *Tricholoma matsutake* was ≥10 m [45–48]. The histgram of genetic distance among sporocarps in Group II (identified as *H. cylindrosporum*) was also bimodal (Fig. 1C). However, only a couple of sporocarps were included in the fewer SNPs peak, suggesting that these sporocarps were clones connected by underground mycelium, and the other sporocarps might belong to genetically distant multiple small mycelia. Dahlberg and Stenlid [45] reported that the majority of *S. bovinus* mycelia in a pine-birch stand were small and produced only a single sporocarp, although some of the mycelia grew large (≥20 m) enough to produce dozens of sporocarps.

The directionality of information flow across sporocarps [9] was also observed in the present study. During the resting period 1, sporocarps were categorized into two types based on the information flow pattern regardless of the species. One consists of sporocarps with strong (large *ETE*) and quick (large *tp*) information flow when they are “causal” (sender). The other is sporocarps with strong and quick information flow when they are “result” (receiver). In addition, a positive correlation between *ETE* and *tp* suggested that strong information flows are transmitted quickly. This categorization was not stable and changed temporarily. However, it was not clear what factors influenced the flow strength and direction and how these fluctuated temporarily. If the electrical potential was transmitted through an underground mycelial network, the spatial location of sporocarps within the mycelial network topology could affect the information flow magnitude and direction, which may fluctuate due to environmental conditions and the physiology of mycelium. Visualized material flow across the mycelial network revealed that the flow direction was outward in the case of growing mycelium, whereas reciprocal transmission was observed in mature mycelium [49–51].

Interestingly, the effect of artificial stimulation on information flow across sporocarps differed depending on the stimulation type. The addition of tap water to the base of sporocarp#1 significantly increased the electrical potential of sporocarp#1 and the average *ETE* among sporocarps within 30 min in both trials. As previously reported, the electrical potential of wild sporocarps increases after rain [9]. Similarly, the addition of tap water to sporocarps and mycelium increased their electrical potential [52]. Water is important to retain physiological processes within cells and produce turgor pressure, which is necessary for hyphal tip growth [2]. Therefore, spatially heterogeneous distribution of water may be important information for fungi, and thus the addition of water to a particular site could induce information flow across the mycelial network. Fukasawa et al. [53] reported that the addition of sterilized wood bait to a certain position of a growing mycelium of the wood decay fungus *Pholiota brunnescens* promoted the electrical information flow from the bait position to other parts of the mycelium. Given that absorbed water would be transmitted through hyphae and improve the cells’ activity at the destination [54,55], this water flow and subsequent activation of forwarded sporocarps could be detected as information flow in this system. In contrast, watering the entire area substantially reduced information flow across the sporocarps. Although we require replicates of this treatment to confirm its reproducibility, the reduction of information flow was significant and worthy of discussion. As opposed to adding water to just one sporocarp, water application to all the sporocarps may have activated all the sporocarps independently, making the patterns of electrical activity less related to each other.

Among the emerged sporocarps in this study, *H. danicum* is known as an ammonia fungus [56] and is expected to be activated by the presence of ammonia. However, *H. cylindrosporum* is known to exhibit reduced growth with ammonia compared with that of other nitrogen sources such as nitrate [57]. The addition of urine as an ammonia precursor had little effect on information flow among the sporocarps. This might be due to the slow process of urine hydrolysis to ammonia. Approximately five days are required at <20°C to alter the soil microbial community and process the urea in urine to ammonia [58]. Considering the lower temperature of the recording period (late October, mean 6.6– 12.3°C), it may require longer. Therefore, the time window to detect information flow (30 s) and the whole recording period of electrical potential (3 days) in the present study might be too short to detect the physiological change of fungi related to urine hydrolysis. As shown by Fukasawa et al. [53], recording for a longer time (≥100 days) may be required to detect a slow process. However, recording the electricity of wild sporocarps using electrodes for a long time is challenging because of the following reasons. First, the short longevity of soft sporocarps of *Hebeloma* spp. Putting electrodes onto underground hyphae may be a solution, but it is challenging because of the small size of hyphae. Second, fluctuation of multiple factors over a longer time in the field makes it challenging to detect the effect of ammonia. The simultaneous recording of environmental factors, such as air temperature, precipitation, light, soil temperature, and soil moisture, and including these in the analysis, might be a solution to an extent.

To summarize, the findings of this study imply that fungal sporocarps in the field may exchange information intra- and inter-specifically in response to environmental conditions. To verify whether the detected information flow constitutes functional signaling, physiological or functional changes in the recipient sporocarps must be examined. As sporocarps are reproductive organs for spore dispersal, they may exhibit lower electrical activity than the vegetative mycelium responsible for growth and nutrient uptake [59]. Future efforts should develop methods for measuring the electrical potential of the subterranean mycelium to elucidate the patterns and mechanisms of fungal electrical communication.

## Author contributions

YF conceptuarize the study and designed the experiment; DA and TT measured the electricity; DT conducted molecular analysis; DA and YO conducted causality of electricity data; YF and DT wrote the manuscript with imput from all authors; YF, DT, and YO reviewed and edited the manuscript. All authors approved the final version.

## Data aailability

The sequences related to fungal sporocarps have been uploaded to the NCBI database (accession#: LC844115 for Group I; LC844116 for Group II). All data are available from correnponding author on personal request.

