## Supplementary material for "Electrical information flows across the sporocarps of two ectomycorrhizal fungi in the field": ElectricMushroom2_SuppleFig.pdf

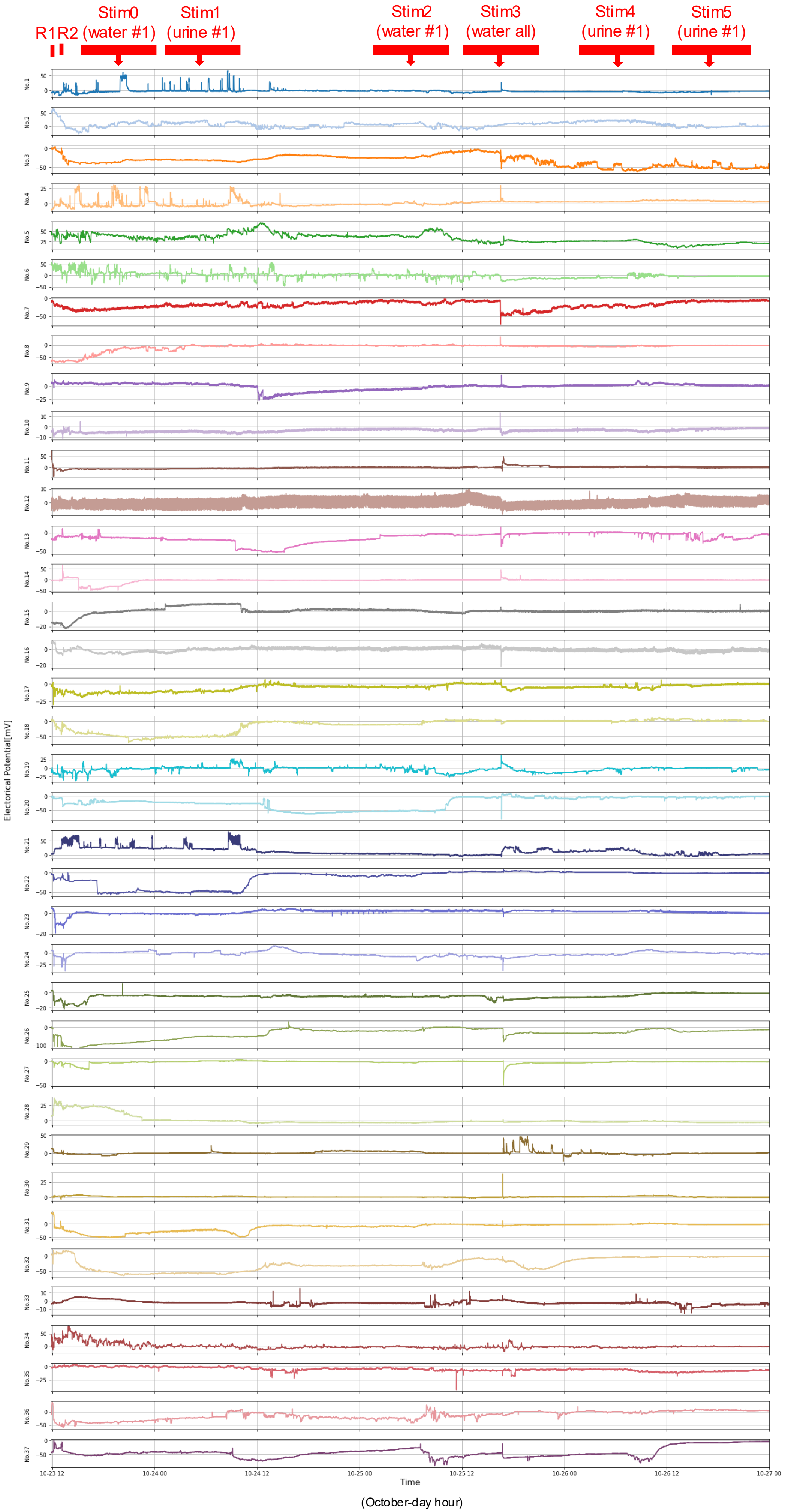


Fig. S1 The full data of 37 sporocarp's electrical potentials and time windows used for causality analysis (red bars).

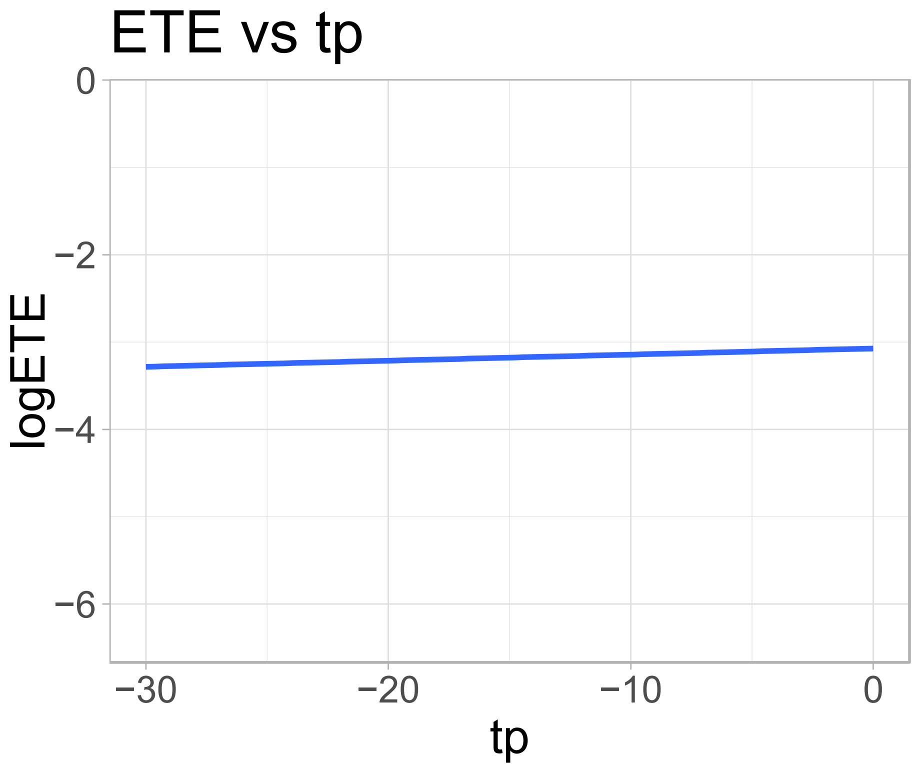


Fig. S2 Correlation between *tp* and *ETE* (logarithm) during the resting period 1. Pearson's $r = 0.0538$; $t = 1.9649$; $df = 1330$; $p = 0.04963$.
